# The biology of aging in a social world: insights from free-ranging rhesus macaques

**DOI:** 10.1101/2023.01.28.525893

**Authors:** Laura E. Newman, Camille Testard, Alex R. DeCasien, Kenneth L. Chiou, Marina M. Watowich, Mareike C. Janiak, Melissa A. Pavez-Fox, Mitchell R. Sanchez Rosado, Eve B. Cooper, Christina E. Costa, Rachel M. Petersen, Michael J. Montague, Michael L. Platt, Lauren J.N. Brent, Noah Snyder-Mackler, James P. Higham

## Abstract

Social adversity can increase the age-associated risk of disease and death, yet the biological mechanisms that link social adversities to aging remain poorly understood. Long-term naturalistic studies of nonhuman animals are crucial for integrating observations of social behavior throughout an individual’s life with detailed anatomical, physiological, and molecular measurements. Here, we synthesize the body of research from one such naturalistic study system, Cayo Santiago Island, which is home to the world’s longest continuously monitored free-ranging population of rhesus macaques. We review recent studies of age-related variation in morphology, gene regulation, microbiome composition, and immune function. We also discuss ecological and social modifiers of age-markers in this population. In particular, we summarize how a major natural disaster, Hurricane Maria, affected rhesus macaque physiology and social structure and highlight the context-dependent and domain-specific nature of aging modifiers. Finally, we conclude by providing directions for future study, on Cayo Santiago and elsewhere, that will further our understanding of aging across different domains and how social adversity modifies aging processes.

## 1. INTRODUCTION

The population of older individuals worldwide is growing at a rapid pace. By 2030, one in six individuals will be aged 60 years or older, and by 2050 this proportion is expected to be one in three (World Health Organization, 2022). With this aging population comes an increased societal impact of diseases of aging, including numerous cardiovascular, immunological, and neurodegenerative diseases (Niccoli and Partridge, 2012). Understanding the multi-faceted nature of aging, and how the environment modifies these processes, is critical for successfully navigating the impending demographic shift. To do so, we must identify the factors that can extend disease-free years of life–known as the “healthspan”. One factor that impacts healthspan is the social environment, which can reduce disease risk and improve survival outcomes (Holt-Lunstad et al., 2010). According to one meta-analysis of over 70 human studies, loneliness and social isolation are associated with a 26% and 29% increased likelihood of mortality, respectively (Holt-Lunstad et al., 2015). Across social mammals, the most socially-integrated individuals live longer and healthier lives (Campos et al., 2020; Snyder-Mackler et al., 2020), and show delayed aging (Chin et al., 2018; Ertel et al., 2008; Loucks et al., 2006). How the social environment “gets under the skin” to affect health, and more specifically senescence, remains an active area of investigation.

To better understand the biology of aging and its modifiers, we can turn to animal systems that share similar biology and social environments with humans. Here we synthesize multidisciplinary efforts to understand how sociality shapes aging processes in one such system: a free-ranging colony of rhesus macaques living on Cayo Santiago Island, Puerto Rico. We begin by outlining why non-human primates are a valuable model for human sociality and aging. We then provide a brief historical background on the Cayo Santiago field site and detail the breadth of behavioral and biological data generated there over the past ~65 years. We summarize recent studies investigating aging and social modifiers of aging through a lens that integrates behavioral, genomic, neuroanatomical, gastrointestinal, immunological, hormonal, and morphological data. Finally, we outline how these findings can be combined to build a holistic understanding of the social modifiers of aging in a naturalistic primate population.

### 1.1 Non-human primates as a key model for human sociality and aging

Questions regarding the interplay between environment and health can be impractical to study in humans for many reasons, including exceptionally long lives, limitations and bias in self report measures, difficulty obtaining repeated measures across the lifespan, and concerns about consent and confidentiality. Further, variation in the human social environment is often confounded by other variables, such as variation in diet and access to healthcare. Studying non-human animals has the potential to circumnavigate these issues. Aging studies in short-lived organisms have laid the essential groundwork in the field (Tsurumi and Li, 2020; Yuan et al., 2011; Zhang et al., 2020), but longer-living species that share a more recent evolutionary history with humans may more readily translate to human biology. This positions non-human primates as critical models in which to study aging and how the social environment affects it (Colman, 2018). Indeed, several studies have begun to address the role of sociality in health and aging, particularly among wild primate populations where social behaviors are not constrained by laboratory settings and health is not managed through veterinary interventions (Archie et al., 2014; Campos et al., 2020; Lehmann et al., 2016; McFarland and Majolo, 2013; Silk et al., 2010).

Rhesus macaques are a particularly useful non-human primate model owing to their frequent use in biomedical research since the late 1800s (Cooper et al., 2022a). Laboratory and observational studies have demonstrated shared genetics, anatomy, neurology, physiology, immunology, behavior, and life-history with humans (Chiou et al., 2020; Colman, 2018; Roth et al., 2004). In particular, like humans, rhesus macaques form relationships based on positive interactions with kin and non-kin, which predict their survival (Brent et al., 2013; Ellis et al., 2019). They also form hierarchically organized societies in which social status can be reliably quantified through measures of dominance rank, that is, an individual’s position in the social hierarchy (Thierry et al., 2004). High status can confer health benefits while low status can result in poor health and fitness outcomes (Blomquist et al., 2011; Michopoulos et al., 2012; Snyder-Mackler et al., 2016; Tung et al., 2012). Additionally, rhesus macaques display developmental and aging trajectories similar to humans, but at a rate of about three to four times faster (Figure 1) (Chiou et al., 2020; Colman, 2018). Overall, these similarities make rhesus macaques an exceptionally well-suited animal model for examining social determinants of health and aging that may translate to humans.

**Figure 1.**
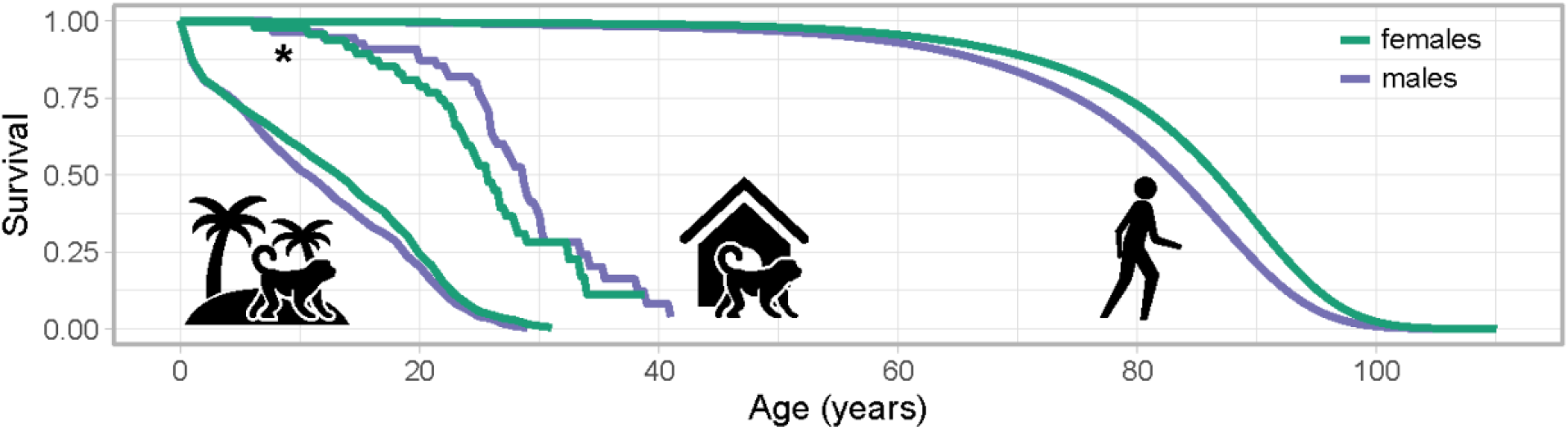
Aging in macaques and humans. Rhesus macaques display a similar but condensed survival trajectory compared to humans, aging at a rate of about three to four times higher. Median lifespan for captive rhesus macaques (indicated by an asterisk) is likely lower than indicated here, as the survival curve is generated from a previously published study (Mattison et al., 2017) that enrolled monkeys cross-sectionally, potentially introducing selection bias and minimizing sources of infant mortality. Reprinted from “Rhesus macaques as a tractable physiological model of human ageing,” by KL Chiou et al., 2020, *Philosophical Transactions of the Royal Society B: Biological Sciences, 375*. Copyright 2020 by The Royal Society. Reprinted with permission.

### 1.2 The deeply-phenotyped Cayo Santiago macaques

Cayo Santiago, a 15.2-hectare tropical island off the southeast coast of Puerto Rico, is home today to approximately 1,500 rhesus macaques. This field site, which is managed by the Caribbean Primate Research Center (CPRC) at the University of Puerto Rico, is the longest-running and most intensively studied population of nonhuman primates in the world (Cooper et al., 2022a). All resident rhesus macaques on the island are the descendants of 409 animals introduced to the island from India in 1938 by primatologist Clarence Ray Carpenter for behavioral research. The macaques are provisioned with food and water, but otherwise receive minimal intervention and manipulation: they are free to form social groups and mating pairs as they please. The monkeys do not have any natural predators on the island. Lifespan on the island is significantly shorter than it would be in captivity; females have a median lifespan of 18 years and a maximum lifespan of 31 years (Brent et al., 2017) (Figure 1). Importantly, the combination of exceptionally high population density (Cayo Santiago: 1,500/15ha, New York City in 2000: ~1,600/15ha, (US Census Bureau, 2001)), food provisioning, and absence of natural predators makes the Cayo Santiago macaque population a model for human urban society, where competition for resources and pressures for survival mainly derive from conspecifics.

One of the biggest assets of the Cayo Santiago field site is the feasibility of collecting both longitudinal and cross-sectional data. Continuous longitudinal data collection on Cayo Santiago was initiated by the CPRC in 1956 to monitor births, deaths, and changes in social group membership. Since the late 1950’s, researchers have independently collected data sporadically over consecutive months or years, including behavioral data, urine, feces, and blood samples. More recently, a multi-institute collaborative team interested in quantifying heterogeneity in aging trajectories has begun collecting longitudinal data on the Cayo Santiago rhesus macaques. This team, the Cayo Biobanking Research Unit (CBRU), has begun routinely collecting a broader set of noninvasive longitudinal measures (e.g., behavioral data, gait speed, urine, and fecal samples) year-round, as well as a standardized set of physiological measures (e.g., blood, soft tissue morphometrics, eye exams) during yearly capture-and-release sampling of a subset (N≈100 per year) of individuals on the island over the past decade (Figure 2).

**Figure 2.**
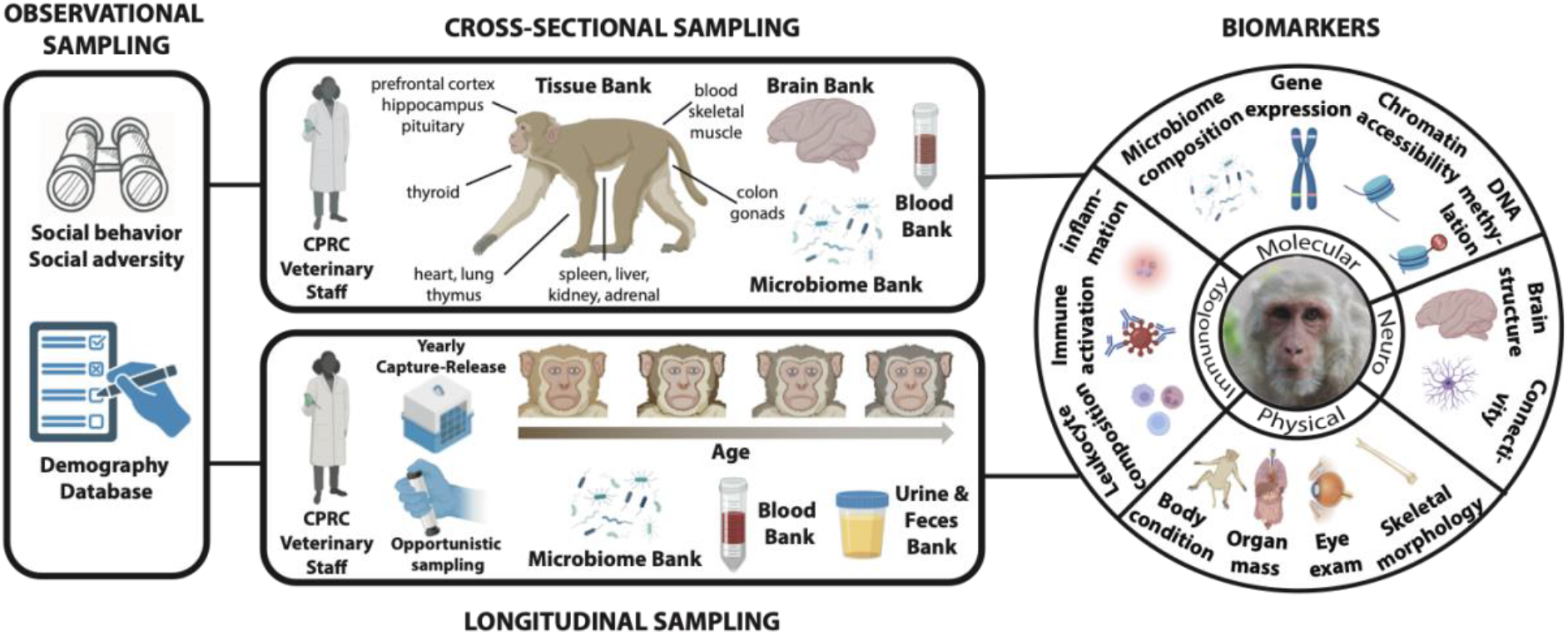
Deeply phenotyped Cayo Santiago rhesus macaques. Demographic data collection (i.e., births, deaths, group membership) on Cayo Santiago was initiated in 1956. Since then, data collection methods have continued to expand: biological samples and behavioral data are collected both cross-sectionally and longitudinally across hundreds of individuals. From these biological samples, it is possible to generate numerous biomarkers of aging. This combination of rich biological, behavioral, and demographic data allows an unparalleled phenotyping and characterization of sociality and aging across biological systems in this non-human primate population. Original macaque photo by L.J.N. Brent. All other graphics were created with BioRender.com.

In addition to these longitudinal measures, cross-sectional samples have also been collected from the Cayo Santiago rhesus macaques. The CPRC has collected complete skeletal remains from deceased individuals since 1971 and blood samples used for assigning genetic parentage since 1992. Cross-sectional samples have also been collected by researchers from various institutions, including blood samples used for a variety of genomic, immunological, and hormonal analyses (Widdig et al., 2016) and cognitive measures (Drayton and Santos, 2016). Due to food provisioning and the absence of natural predators, the average annual growth rate of the Cayo Santiago macaque population exceeds that of wild groups, making population control necessary through capture and removal of animals (Hernandez-Pacheco et al., 2016). Starting in 2016, the CBRU has collected and cataloged a plethora of samples from animals removed from the island, including, but not limited to, postmortem tissues from every major organ (e.g., brain, heart), swabs for microbiome analyses, blood, primary cell cultures, skeletons, and soft tissue measurements (Figure 2). Crucially, these samples are collected from socially phenotyped monkeys of known age and sex, enabling integration of varied axes of information.

The unique breadth and depth of data, both longitudinal and cross-sectional, collected from the Cayo Santiago rhesus macaques has allowed detailed exploration of age-related variation across multiple biological domains (e.g., musculoskeletal, immunological, molecular) as well as the social modifiers of aging. Recent efforts led by the CPRC and CBRU have greatly expanded the types of biological data collected and refined the systematic collection of behavioral data to address questions surrounding social adversity and aging. Thus, a majority of the studies discussed in this review are projects conducted by members of the CBRU collaborative team. However, relevant work conducted by other researchers utilizing the Cayo Santiago field site is also included where appropriate.

## 2. AGING IN THE CAYO SANTIAGO RHESUS MACAQUES: A MULTIFACETED PHENOMENON

A key question in biology is: To what extent is age reflected in different biological domains? Further, a thorough characterization of age across domains is necessary for exploring how age may be modified by the social environment. Below, we review recent findings focused on age-related variation in morphology, gene regulation in the brain, microbiome composition, and immune system function. Together, these results are paving the way towards an integrated understanding of how biological systems vary with age.

### 2.1 Morphology

Our bodies undergo numerous morphological changes as we age, including alterations to musculoskeletal and ocular function. Musculoskeletal diseases are one of the leading contributors to global disease burden in older individuals (Prince et al., 2015) and three of the four main causes of blindness are strongly associated with age (Steinmetz et al., 2021). However, there remains a need to explore the relationship between aging and morphological changes in large populations without major confounding variables such as diet or exercise.

The detailed skeletal collection and cross-sectional measurements of mobility and body mass collected on the Cayo Santiago rhesus macaques have enabled researchers to probe how the skeletomuscular condition of older individuals differs from that of their younger conspecifics. Results from this population demonstrate that, similar to humans, older individuals display significantly less joint mobility than younger individuals and present evidence of degenerative joint disease, osteoporosis, osteopenia, and arthritis (Cerroni et al., 2000; Kessler et al., 1986; Renlund et al., 1986; Turnquist and Kessler, 1989). In females, bone mineral density and bone mineral content steadily decline after 17 years of age (Cerroni et al., 2000). Notably, age-related variation in body mass remains unresolved, with some evidence suggesting older females display lower body mass than younger females (Hoffman et al., 2010) while other findings indicate body mass does not differ between adulthood and late life in either sex (Turcotte et al., 2022). Differences in methodological approach may explain these inconsistencies and future work will be necessary to confirm whether age-related variation in body mass exists in this population.

The onset of bone diseases such as osteoporosis coincides with reproductive decline in this population. Female rhesus macaques display reproductive senescence characterized by a rapid decline in fertility after age 17 (Lee et al., 2021). Given that a steady decline in bone mineral density also begins at age 17, this change in bone condition may be akin to the accelerated rate of bone loss seen in perimenopausal women (Lo et al., 2011). Older males on Cayo Santiago also experience bone diseases that may be related to reproductive decline, as high-ranking males over the age of 18 sire fewer offspring than younger conspecifics, unrelated to mating effort (Milich et al., 2020). Older males also display lower fecal androgen levels than younger males during the birth season (Higham et al., 2013), and future research in the population will be useful for determining whether hormonal changes related to reproductive senescence are responsible for the skeletomuscular changes observed in older individuals.

The array of data types collected from the Cayo Santigo rhesus macaques makes it possible to evaluate morphological changes in domains other than the skeletomuscular and reproductive systems. Recent research on the island has focused on age-related variation in ocular morphology. Rhesus macaques are particularly well suited for eye-related studies since they share several ocular features with humans that are absent in classic rodent and rabbit models. Eye exams demonstrate older macaques on Cayo have thinner corneas, greater axial length, and less myopic shift compared to younger individuals, indicative of age-related ocular variation that is also observed in humans (Fernandes et al., 2022; Galgauskas et al., 2013; McCullough et al., 2020; Verkicharla et al., 2020). Incidence of cataracts and retinal drusen are also significantly higher in older individuals than younger conspecifics (Fernandes et al., 2022). Longitudinal studies will be necessary to determine whether these changes truly represent markers of aging rather than arising from population-level factors such as differential mortality (i.e., selective disappearance) (Nussey et al., 2008). Similar to humans, retinal cone gene expression in this population does not show any significant age-associated differences (Munds et al., 2022) and additional research combining cross-sectional ocular measures with transcriptomic data from the eye may provide novel insights into the genetic underpinnings of age-related eye diseases.

### 2.2 Gene Regulation in the Brain

In addition to morphological differences observed between young and old individuals, age-associated declines in brain function are well established in humans and other animals (Mattson and Arumugam, 2018). However, our understanding of how the brain varies with age, especially at the level of gene regulation, is far from complete. This is in part due to the reliance on post-mortem brain tissue samples from older or diseased individuals since collecting brain tissue from healthy humans is generally infeasible. In the Cayo Santiago study system, cross-sectional sampling of tissues from rhesus macaques removed as part of population management circumvents these limitations. Recent work profiling the transcriptomes of 15 brain regions acquired from healthy adult Cayo Santiago rhesus macaques revealed numerous brain-region specific and multi-region signatures of age (Chiou, Decasien et al., 2022). Many of these multi-region (i.e., “whole-brain”) age-associated genes were also identified in humans as markers of a diseased state in Alzheimer’s disease. Since the macaques in this study did not exhibit any pathologies associated with Alzheimer’s disease, this suggests that some transcriptomic hallmarks of Alzheimer’s disease may manifest earlier in the course of aging, even before neurodegenerative pathologies are present. Further, the accuracy of correctly predicting the sex of these individuals based on non-Y chromosome genes decreases with advancing age– particularly in males (DeCasien, Chiou et al., 2022). This suggests that age-associated breakdowns in tightly regulated suites of genes (transcriptional “decoherence”) may be exacerbated in the brains of males compared to females. Overall, these results constitute the most comprehensive transcriptional analysis of age in the brain in a free-ranging non-human primate and demonstrate age-associated molecular markers may precede physical signs of neurological disease. Future integration of this transcriptomic data with matched neuroanatomical phenotypes (Testard et al., 2022) will be invaluable for identifying the relationship between molecular markers of age and morphological variation, including pathophysiological signs of neurodegenerative diseases.

### 2.3 Microbiome Composition

Recent advances have elucidated the extensive bidirectional signaling network between the brain and the microbiota that reside in the body. For example, age-related changes in the composition of microbial communities in the gastrointestinal tract have been suggested to play a role in the development of dementia and Alzheimer’s disease and to modulate cognitive function, immunity, and frailty (Askarova et al., 2020; Britton and McLaughlin, 2013; Claesson et al., 2012; O’Toole and Jeffery, 2015; Zapata and Quagliarello, 2015). Cross-sectional microbiome samples from different areas of the body as well as longitudinal collection of fecal samples from the Cayo Santiago rhesus macaques are particularly useful for advancing our understanding of age-related microbiome changes and their relationship to health. Indeed, quantifying microbiome variation across a large sample of Cayo Santiago rhesus macaques spanning infancy to old age has revealed significant developmental patterns in the gut, oral, and penile microbiomes (Janiak et al., 2021). The penile microbiome is relatively variable across age groups, and strong age-related differences observed between 1-4 year-olds and other age groups may be related to the onset of sexual behaviors. In contrast, there were no detectable age-related associations in the vaginal microbiome. Furthermore, infant gut and oral microbial communities are relatively distinct from all other age groups. The infant gut has significantly higher relative abundances of milk-digesting genera (e.g., *Bifidobacterium* and *Bacteroides*), and lower abundances of fiber-digesters (e.g., *Ruminococcus*, *Fibrobacter*, and *Treponema*) compared to older individuals. Notably, the gut and oral microbiota of adult macaques remains relatively stable, including into advanced age, contrary to findings in older humans (Wilmanski et al., 2021). A potential explanation for this discrepancy is that age-related microbiome variation found in humans may not be due to the aging process itself, but rather due to lifestyle and dietary changes or various medical interventions that are a common feature of aging in modern-day humans. Future longitudinal sampling of microbial communities (e.g., fecal, skin, armpit) in the Cayo Santiago macaques will elucidate the stability of the microbiota within individuals while cross-sectional samples of microbial communities in the gastrointestinal tract collected at the same time as brain biopsies will provide insight into the relationship between the brain and gut microbiome.

### 2.4 Immune System Function

The immune system interacts with almost every facet of our biology, including the skeletomuscular system, brain, and gut microbiome. Immune systems in older individuals are typified by chronic, low-grade inflammation, which contributes to higher incidence of inflammatory diseases and is associated with greater susceptibility to cancers and infection (Shaw et al., 2010; Weyand and Goronzy, 2016). Characterizing age-associated variation in immune functioning is essential to understanding the biological mechanisms underlying immunological aging. In the Cayo Santiago system, blood samples are routinely collected both longitudinally for the same individuals across several years and cross-sectionally among the population, allowing for a fine-grained investigation of immune activation and age. Analysis of these blood samples indicates that transcriptomic age-variation in rhesus macaques mirrors that of humans (Chiou et al., 2020). Additionally, non-invasive measures (e.g., urine) are collected longitudinally from the same individuals with even greater frequency than blood samples, enabling regular monitoring of immune activity.

A number of age-associated immune differences have been documented from blood and urine samples from the Cayo Santiago population, suggesting immune dysregulation with age. Blood transcriptomic analyses indicate that genes associated with inflammation and innate immune activity are expressed at higher levels in older individuals (termed “inflammaging”) (Chiou et al., 2020; Watowich et al., 2022). Further, older individuals have higher proportions and activation of innate immune cell populations, and a lower proportion of adaptive (e.g., anti-inflammatory) immune cells in the blood (Rosado et al., 2021; Watowich et al., 2022). Older females also show higher levels of plasma interleukin-1 receptor antagonist than younger conspecifics (Hoffman et al., 2011). Together, these findings suggest increased age is associated with greater inflammatory pathway activation and a greater proportion of inflammatory immune cells in the blood. Importantly, many of these immunological differences can be detected non-invasively. For example, urinary neopterin, a biomarker of innate immune system activation in primates, increases with age in rhesus macaques (Cooper et al., 2022b). As such, urinary neopterin holds promise as a predictive biomarker of age-related health declines associated with immune system functioning. Combining invasive and non-invasive measures from the same individual provides the unique opportunity to validate non-invasive measurements in the field, which can then be used by other scientists studying wild populations where invasive methods are not possible.

Overall, age is associated with concurrent variation across multiple biological systems in the Cayo Santiago rhesus macaques. There is, however, substantial inter-individual variation in aging phenotypes in this population, with some individuals exhibiting an older or younger phenotype than expected for their chronological age (i.e., time since birth). Environmental stressors such as resource scarcity, on the one hand, and social support on the other, are major determinants of senescence trajectories (Holt-Lunstad et al., 2015, 2010; Snyder-Mackler et al., 2020). With over a decade of longitudinal social behavioral data, natural variation in environmental stressors, and a wealth of biological sampling, Cayo Santiago provides a unique opportunity to investigate the socio-ecological determinants of health and aging.

## 3. ECOLOGICAL MODIFIERS OF AGING

A tremendous source of instability for humans and other animals is exposure to natural disasters, such as earthquakes and tsunamis, and massive weather events such as hurricanes. Extreme weather can cause widespread destruction of the natural landscape, resources, and infrastructure, all of which can profoundly disrupt the lives of humans, including a sustained decline in both mental and physiological health (Adams et al., 2011; de Oliveira et al., 2021; Hussain et al., 2011; Kario and Ohashi, 1997; Shultz and Galea, 2017; Thomas et al., 2018). With the intensifying climate crisis, devastating storms are expected to become less predictable and increase in both frequency and force. Therefore, understanding how such events influence aging and disease is critical to predict the effects of climate change on global health.

In 2017, Hurricane Maria, a devastating category 4 storm, hit Cayo Santiago leading to the complete destruction of the infrastructure on the island and a >60% drop in vegetation overnight (Testard et al., 2021). Surprisingly, only 2% of the population died from the hurricane, compared to previously reported death rates of 35-60% following natural disasters in wild animal populations (Testard et al., 2021), likely due to food-provisioning on the island. Hurricane Maria allowed scientists to investigate the physiological effects of an extreme environmental stressor and its impact on health and aging in whole macaque social groups.

Watowich and colleagues (2022) found broad similarities between the effects of age and exposure to Hurricane Maria on peripheral immune cell gene expression. Genes more highly expressed in older individuals and in individuals that experienced the hurricane were enriched in inflammatory pathways and loss of protein homeostasis, suggesting possible pathways by which the environment and age can both lead to immune dysregulation. Moreover, individuals that experienced the hurricane had a gene expression profile that was an average of 1.96 years older than individuals who did not experience the hurricane–corresponding to an increase of approximately 7 to 8 years in a human life (Watowich et al., 2022). While these results were based on a dataset of primarily cross-sectional samples opportunistically taken before and after Hurricane Maria, ongoing longitudinal data collection from the Cayo Santiago macaques will build upon these results to test for the effects of environmental adversity within individuals as they age. Together, this will have great translational relevance for human health as our own species confronts the impacts of climate change.

Exposures to hurricanes early in life can also precipitate lifelong changes that may impact health, reproduction, and survival. On average, female macaques on Cayo Santiago that experienced hurricanes had a delayed reproductive start (i.e., later age at first birth). However, females appeared to compensate for this delay by producing more offspring during their prime reproductive years compared to females who did not experience a hurricane (Luevano et al., 2022). The natural disaster-related changes in reproductive biology may lead or be linked to modified aging trajectories which have yet to be explored.

While Cayo Santiago is very well-suited to study the effects of extreme weather events such as hurricanes on health and aging, it exhibits limited environmental variation given its location close to the equator, food provisioning, and absence of predators. Sources of environmental variation should also be explored at other field stations that feature seasonal variation in temperature, resource distribution, and/or predation.

## 4. SOCIAL MODIFIERS OF AGING IN A NATURALISTIC PRIMATE SOCIETY

When individuals face environmental adversity, such as a major hurricane, they may rely on their social support network to mitigate some of the ensuing negative consequences–called “social buffering” (Cohen and Wills, 1985). Social support can be conferred by a few strong connections to key partners, or by numerous weak connections that broadly connect individuals to their network (Ellis et al., 2019). Social status is another, sometimes orthogonal, dimension of sociality which confers advantages such as competitive access to resources. We examine how different dimensions of sociality modify aging trajectories at two levels, ultimate and proximate. At the ultimate level: What advantages do macaques’ social relationships provide in terms of survival and reproduction? At the proximate level: How does sociality “get under the skin” to shape biological function and health? We then discuss the extent to which the relationships between sociality and health are context-dependent.

### 4.1 Ultimate pathways through which sociality affects health and aging

Long-term monitoring of the Cayo Santiago macaque population has provided evidence of the importance of social relationships and social status in helping individuals cope with natural disasters, conspecific competition, and disease exposure, all of which are known causes of early death (Alberts, 2019; Pavelka et al., 2007; Pavez Fox et al., 2022; Pavez-Fox et al., 2022).

In the aftermath of Hurricane Maria, individuals became more tolerant of each other (400-600% increase in the time spent in proximity to other monkeys) and built social connections with new, unfamiliar partners instead of reinforcing existing relationships (Testard et al., 2021). Increased social tolerance may be a response to the loss of vegetation which dramatically reduced the shady surface areas which monkeys could use to regulate body temperature in the excruciating Caribbean heat. Widening of one’s social network may have increased access points to now scarce and valuable shady spots on the island. Better access to shade leads to lower heat stress which, compounded over the years, can putatively lead to better health and aging trajectories, though future work is necessary to confirm this hypothesis.

Cayo Santiago macaques are food-provisioned, and their home island is devoid of predators. Thus, outside of major destructive weather events, the pressures for survival mainly come from conspecifics. Injuries, generally inflicted by conspecifics during aggressive interactions, are a major cause of death in this population (Pavez-Fox et al., 2022). Indeed, injured adults are 3 times more likely to die in ensuing months than non-injured counterparts. Importantly, injury-related death may be mitigated through social capital. Across 1601 individuals, both matrilineal rank and the number of relatives in one’s group (proxies for social status and social connectedness in females, respectively) predicted the likelihood of getting injured. Having more relatives in a group likely results in having more partners available to provide agonistic support (Silk, 1982) or more opportunities to access resources without engaging in aggressive encounters (Samuni et al., 2018). In contrast, once injured, social resources have no impact on rhesus macaque survival, suggesting hygiene-related behaviors, such as grooming, might not aid in recovery (Pavez-Fox et al., 2022).

In addition to conspecific-induced injury risks, parasites and infectious diseases constitute major challenges associated with group-living. Social contact and shared space can increase disease transmission, while individual differences in social resources can help prevent infections. For instance, high social status may reduce susceptibility to infections by promoting better health and immunity (Pavez-Fox et al., 2021; Sapolsky, 2005) or by mitigating exposure to parasites (Müller-Klein et al., 2019). In this population, spatial proximity to partners, but not grooming or social status, predicted lower water-borne parasite load (Pavez Fox et al., 2022). These results suggest proximity to other individuals might enable access to better quality or cleaner resources through social tolerance.

As individuals age, their priorities and needs shift, along with their experience of the social world and ensuing social strategies. Humans exhibit age-associated social selectivity, through which individuals prioritize meaningful relationships and narrow their social networks to key (or “preferred”) individuals as they age, rather than maintaining broad social networks with weaker connections. For the Cayo Santiago rhesus macaques, the importance of specific social connections for health also changes across the lifecourse: adult females with more relatives in their group lived longer, but this was not the case for old females (Brent et al., 2017). Further, increased social selectivity with age has also been observed in female Cayo Santiago rhesus macaques (Siracusa et al., 2022), suggesting conserved patterns of social aging in humans are deeply rooted in primate evolution and have adaptive value. The causes and consequences of these changes in sociality with age are still unknown and may be an important source of unexplained variation in individual patterns of senescence (Siracusa et al., 2022).

In sum, social relationships can provide better access to scarce resources and support during competition. Moreover, the quality and quantity of social relationships are associated with differences in individuals’ biology. In the next section we review the proximate mechanisms through which sociality improves health and aging trajectories.

### 4.2 Proximate mechanisms: Biological underpinnings of the social determinants of health

A large body of work highlights the immune system as a key mechanism by which sociality “gets under the skin” to affect health. In humans, social support (i.e., number and strength of social connections) and social status (e.g., socio-economic status, resources) strongly predict immune system activation, such that individuals with more social connections and higher social status generally have more favorable immune outcomes (Holt-Lunstad et al., 2015; Uchino et al., 2018). In rhesus macaques, experimental manipulation of social status in females causally alters immune cell gene regulation. Low-status females show signatures of heightened inflammation overall and in response to simulated infections *in vitro*, putatively in response to chronic stress and, ultimately, glucocorticoid resistance (Snyder-Mackler et al., 2019, 2016; Tung et al., 2012). This relationship between sociality and immunity has also been probed in Cayo Santiago’s macaque population, which suffers from fewer confounding variables than humans (e.g., diet and exercise) and benefits from natural variation in sociality that is nonexistent in laboratory macaques. Macaques of all ages and sexes with more grooming partners have lower white blood cell counts (Pavez-Fox et al., 2021), suggesting lower systemic inflammation and reduced susceptibility to pathologies (Karthikeyan and Lip, 2006; Sabatine et al., 2002). In addition, social status has a strong effect on the proportion of specific immune cell types (e.g., natural killer cells and T cells) in both male and females rhesus macaques, and higher ranking males have a more pro-inflammatory blood immune cell composition reflecting a stronger innate immune response than lower ranking males (Georgiev et al., 2015; Rosado et al., 2021). Interestingly, the opposite is true for females; low-status females have a more pro-inflammatory immune cell composition than high-status female counterparts (Rosado et al., 2021). Notably, Pavez-Fox and colleagues (2021) did not find a statistically significant association between social status and gross immunity markers such as spleen and liver weights and white blood cell counts, indicating only more fine-grained measures of immune function like immune cell composition index the effects of social status on immunity (Rosado et al., 2021). Together, these findings suggest that social integration may promote anti-inflammatory immune phenotypes in macaques, while the relationship between social status and immune activity is more nuanced.

These findings are consistent with the social ecology of this species in which females maintain a stable social hierarchy throughout their lifespans while male social hierarchies are far more variable because they result from queuing and aggression. The inflammatory profile in low-status females may reflect chronic stress due to social harassment, social isolation or reduced food intake (Sapolsky, 2005; Snyder-Mackler et al., 2019); while in high-status males it may reflect a response to conflict (e.g., injury recovery). Additionally, immune activity may vary over shorter time scales, such as in relation to seasonal behavioral changes, which are only possible to measure with fine-grained repeated sampling. For example, levels of urinary neopterin, an innate immune system activation marker, were also higher in males who spent more time grooming, but not copulating, during the mating season (Petersen et al., 2021). Overall, the links between sociality, immunity, and health are highly nuanced and context-dependent. Future work should account for and further explore these important contingencies.

The brain orchestrates all biological systems within the body while sitting at the interface of the external socio-ecological world and internal regulatory processes. This positions the central nervous system as key to understanding the biological translation of the physical and social environment into health and aging. With respect to the social environment, the brain controls the perception of social cues, consolidates and integrates memories of past interactions, and ultimately uses that information to guide social interactions (Ong et al., 2020; Shepherd and Freiwald, 2018). Primates also track third party interactions to inform understanding of the social hierarchy (Cheney et al., 1986; Cheney and Seyfarth, 2018). The brain then generates appropriate behavioral and physiological responses, including immune activation.

The essential role of the brain in connecting social support and status to biological success is increasingly well understood in invertebrates and model organisms like fish and mice, but remains more opaque in humans and other primates (Utevsky and Platt, 2014). Testard and colleagues (2022) found that in adult rhesus macaques on Cayo Santiago, the number of grooming partners predicted the volume of the mid–superior temporal sulcus and ventral-dysgranular insula, which are implicated in social decision-making and empathy, respectively. Notably, brain structure varied specifically with the number of affiliative social connections, but not other key dimensions of sociality such as social status or position in the social network, as was also the case for certain immune markers (Pavez-Fox et al., 2021). Although an effect of social status could not be detected at a gross neuroanatomical level (Testard et al., 2022), patterns of gene expression in the brain varied with social status and age in a parallel fashion (Chiou, Decasien et al., 2022). Specifically, high-ranking females exhibited younger transcriptomic ages across the brain. Given that aging trajectories may be modified by social experiences, future work on this population may help tease apart the mechanisms through which social experience accelerates or slows aging in the brain.

### 4.3 There are many ways to ‘be social’

In different socio-ecological contexts and at different ages, distinct facets of social behavior might be more important for survival. In female Cayo Santiago rhesus macaques studied before Hurricane Maria, only the strength of connection to favored partners and broad connectivity to the group (number of weak connections) predicted survival, but not indirect connectedness (e.g., number of friends of a friend) and time spent interacting (Ellis et al., 2019). Having more relatives in the group was particularly important for survival in young adult females (Brent et al., 2017). By contrast, after Hurricane Maria, broad connections to the wider social group were more important than investing in existing key relationships such as with kin (Testard et al., 2021). Social status did not predict gross brain structure (Testard et al., 2022) but did predict brain gene expression profiles (Chiou, Decasien et al., 2022). These examples from the Cayo Santiago population demonstrate that, depending on the context, needs, and biological system, certain dimensions of sociality might be more important than others in predicting biological variation and survival. Only large populations with natural variation in sociality, such as the Cayo Santiago macaque population, can facilitate the dissection of context-dependent biological pathways between sociality, aging, and health.

Overall, the Cayo Santiago rhesus macaque population has offered unique and unparalleled insights into the physiological correlates of age and how socio-ecological factors may influence senescence. Next, we propose future avenues of investigation to answer fundamental questions about the relationship between health and the social environment as we age.

## 5. COMBINING EVIDENCE FROM MULTIPLE BIOLOGICAL SYSTEMS

### 5.1 How do within-individual aging trajectories change across biological and social domains?

Many studies evaluating age-related differences in physiology and behavior, including several of the studies discussed above, draw on cross-sectional sampling methods. While these methods allow for a diversity of data to be collected, they only enable insight into population-level processes; it is only possible to determine whether younger and older individuals differ from one another. Further, any difference observed cross-sectionally may be the result of differences among cohorts or selective disappearance, rather than aging itself (Nussey et al., 2008; Siracusa et al., 2022; van de Pol and Wright, 2009). For example, if socially isolated individuals are more likely to die because of predation, at the population-level there may appear to be an age-related change in sociality when, in fact, no within-individual change is present (Siracusa et al., 2022). Longitudinal studies are therefore necessary to establish age-related patterns that result from within-individual aging trajectories (Nussey et al., 2008). Only by conducting longitudinal studies is it possible to determine within-individual rates of aging across different biological domains and begin to disentangle what accounts for heterogeneity in rates of aging between individuals.

Within-individual age-related declines have been assessed previously across several factors such as probability of survival, fecundity, body mass, and immune response in wild populations (Nussey et al., 2008). The Cayo Santiago field site provides a unique opportunity to measure longitudinal aging patterns across many more traits than is possible at most other field sites. Currently, longitudinal data collection on the island includes, but is not limited to, behavioral measures, life-history data, eye exams, soft tissue morphology, gait speed, fecal and skin microbiome samples, blood, urine, and feces (Figure 2), with the diversity of measures collected increasing almost every year. Continued evaluation of within-individual changes in these domains across the adult lifespan will generate one of the most comprehensive descriptions of aging to date, which, in turn, will make the data more valuable each year it is collected. Further, this multi-modal longitudinal data collection will establish the temporal relationship between rates of aging across biological domains, such as whether age-related declines in mobility precede changes in sociality. Such information will help elucidate the direct role of physical age-related changes on social behavior. Importantly, there are some limits to the types of data that can be collected longitudinally on these rhesus macaques, and traits such as solid organ aging will probably never be longitudinally measured in this study system. As a result, laboratory set ups that can collect invasive samples (e.g., muscle and endoscopic biopsies) in a non-lethal manner will be necessary to advance our understanding of aging patterns for certain factors. Overall, the breadth of longitudinal measures collected from the Cayo Santiago rhesus macaques will enable detailed exploration of aging trajectories across several biological and social domains and help to elucidate the sources of heterogeneity in aging rates between and within individuals.

### 5.2 Does social adversity mimic aging at the molecular level?

A fundamental gap in the social aging literature is an understanding of the extent to which age and social adversity are associated with overlapping molecular mechanisms and, ultimately, how social adversity influences the aging process. Studies in humans have demonstrated that many of the diseases associated with social adversity are also common diseases of aging (Barth et al., 2010; Dong et al., 2004; Hawkley and Cacioppo, 2004; Holt-Lunstad et al., 2015; Shalev, 2012). In laboratory studies, experiencing social adversity directly results in shortened lifespan, immune dysfunction, increased cellular markers of senescence, and earlier onset of a number of diseases including cardiovascular disease (Bartolomucci, 2007; Kinnally et al., 2019; Lin et al., 2021; Razzoli et al., 2018; Snyder-Mackler et al., 2016). Further, in the blood, transcriptomic signatures of chronic social stress in rhesus macaques are shared with signatures of advancing age in humans (Snyder-Mackler et al., 2014). Previous studies of the Cayo Santiago rhesus macaques have also illustrated that social status and age show overlapping transcriptional variance in the brain and immune cell composition in the blood (Chiou, Decasien et al., 2022; Rosado et al., 2021). However, the extent to which social adversity recapitulates age more broadly, across molecular mechanisms, types of adversity, and different organs, is still unknown.

To answer this question, it is essential to test for similarities between age and social adversity at multiple levels of gene regulation. Methods to investigate effects on gene regulation include analysis of genetic variants, DNA methylation, gene expression, and chromatin accessibility. Initial analyses integrating the independent effects of social adversity and age at each of these levels would clarify to what extent physiological processes associated with age and sociality are shared. Subsequent analyses integrating transcriptomic, epigenomic, and chromatin accessibility data could determine which genomic regions regulate genes whose expression is significantly altered with age or social adversity. Identifying genetic variants at regions that are associated with both age and response to social adversity could provide new insights into how genetic differences confer susceptibility or resilience to social adversity and aging.

Importantly, these molecular mechanisms must be evaluated across the body since molecular changes in one tissue due to age or social adversity may not be the same in other tissues (Schaum et al., 2020; Slieker et al., 2018; Yang et al., 2015). In the same way, the molecular changes resulting from different types of social adversity, such as early life social adversity and resource competition in adulthood as well as the number, degree, and magnitude of adverse events need to be taken into consideration and tested independently for their similarity to age-associated molecular changes. As such, the extensive behavioral and multi-tissue genomic data collected on the Cayo Santiago rhesus macaques makes this population particularly well suited for investigating whether social adversity recapitulates the effects of age. Samples from over 25 different organs in individuals ranging from infancy to old age, including brain and blood samples, can be analyzed for age-associated alterations in genetic effects, methylation, gene expression and chromatin accessibility. Detailed behavioral data from these same individuals, including various types of social adversity experienced throughout life, will enable investigation of molecular changes that are correlated with different forms of adversity. Comparing age-related and social adversity-related molecular mechanisms will shed light on the extent to which these mechanisms are shared, paving the way for future research into social and molecular modifiers of health and disease.

### 5.3 How do social adversity and microbiome composition affect the risk of developing neurodegenerative disorders?

Our rapidly growing population of older individuals carries a corresponding increase in age-related disease burden, which is particularly acute for age-related brain diseases such as Alzheimer’s disease, dementia, Parkinson’s disease, amyotrophic lateral sclerosis and Huntington’s disease. However, there is substantial variability in the onset and severity of neurodegenerative disease, and emerging evidence suggests that susceptibility and mortality risk are impacted by social adversity and dysbiosis of the gut microbiome. For example, social isolation is associated with an increased risk of developing Alzheimer’s disease and other dementias (Drinkwater et al., 2022), and changes in gut microbiome composition characterize multiple neurodegenerative conditions, including Alzheimer’s disease and Parkinson’s disease (Li et al., 2017; Shen et al., 2017; Vogt et al., 2017). In fact, administration of probiotics or gastrointestinal microbiome transplantation from healthy mice into Alzheimer’s disease mouse models improves memory, reduces brain beta-amyloid and tau aggregation, and decreases neuroinflammation, implicating gut microbiota in neurodegenerative disease moderation (Abraham et al., 2019; Kim et al., 2020).

Neuroinflammation is a common pathophysiological mechanism that underlies many neurodegenerative diseases (Hong et al., 2016). Although aging is generally characterized by a progressive increase in neuroinflammation, precisely how social connections and the gut influence the relationship between aging and neuroinflammation remains poorly understood. In addition, the extent to which these relationships may be mediated by peripheral inflammation (which can induce neuroinflammation) remain unclear (Hoogland et al., 2015). These gaps in part reflect limited access to primary tissue samples and difficulties measuring direct objective measures of social behavior in humans. In rodents, molecular signals from the gut affect many physiological systems relevant to human health and experimentally-induced social deficits can be reversed through microbiota modification. However, rodent models are limited by significant differences in the nature and complexity of social behavior, distinct molecular biology and anatomy of the brain, and divergent gastrointestinal tracts in rodents and humans. Recent studies have validated the rhesus macaque as an important nonhuman primate biomedical model that shares important aspects of complex human social behavior (Deaner and Platt, 2003), underlying neural circuitry (Noonan et al., 2017), signatures of aging in the brain transcriptome (Chiou, Decasien et al., 2022), gastrointestinal structure and function (Chivers and Hladik, 1980), and links between social isolation and health (McCowan et al., 2016).

Notably, the relationships between social adversity, gut microbiome composition, and disease observed in humans are likely to be mediated by biological sex. For instance, although females are disproportionately affected by dementia, afflicted females tend to live longer than afflicted males (Sinforiani et al., 2010), which may be in part due to males being more likely to experience social isolation (Umberson et al., 2022). Although biological sex has historically been ignored in many animal model studies (with a bias towards male-only study designs), the inclusion of both males and females in research is now a priority for many funding agencies (e.g., NIH policy on SABV). Given that the Cayo Santiago population includes approximately equal numbers of males and females, and recent work confirms that rhesus macaques and humans exhibit similar sex differences in the brain transcriptome (Decasien, Chiou et al., 2022), this creates a unique opportunity to consider the potential impacts of biological sex on our variables of interest.

The Cayo Santiago population provides a unique opportunity to identify the biological links between social connectedness, the gut microbiome, and age-related patterns of neuro- and peripheral inflammation, while also considering the possible modulating effects of biological sex. In particular, future cross-sectional studies have the potential to integrate measures of social connectedness, gut microbiome content/diversity, neuroinflammation (derived via e.g., brain gene expression or histological measures), and peripheral inflammation (derived via e.g., cytokine concentrations). This holistic data set would allow us to illuminate not only how social connectedness and the gut microbiome interact with each other, but also how these factors moderate the relationship between age and peripheral/neuroinflammation. In addition, we will be able to examine how these factors moderate the relationship between peripheral inflammation and neuroinflammation, which may provide critical insights into our understanding of potential peripheral biomarkers for neurodegenerative disease risk in humans.

## 6. CONCLUSIONS

The Cayo Santiago study system is a unique natural laboratory that integrates the benefits of studying free-ranging primates and their naturalistic behaviors with the ability to collect invasive and non-invasive biological samples. Thus, the rhesus macaques that inhabit this island are an invaluable tool for characterizing biological changes that occur during aging and how social integration, isolation, and status can modify these processes. Previous research on Cayo Santiago has focused on how age is independently related to certain aspects of social behavior, morphology, physiology, and the immune system. More recent studies have begun to illuminate biological mechanisms that are shared between social adversity and advancing age, particularly with regard to brain gene expression and blood immune cell composition. Some of this work has begun to exploit longitudinal data to move beyond differences between individuals of different ages, to what happens to an individual as it ages across its life course. It is our hope that future research on Cayo Santiago will continue to expand on the available measures of sociality and aging to address unanswered questions about the role of social integration and isolation in shaping signatures of aging across biological levels. Such research can provide critical insights into how social adversity “gets under the skin” and will hopefully spark new advances that mitigate the negative age-related effects of social adversity in humans.

## Acknowledgements

The authors would like to thank the Caribbean Primate Research Center and the staff from the Sabana Seca Field Station and the Cayo Santiago Field Station, without whom this research would not be possible.

